# Liver transplant rejection features memory cell priming, spatially differentiated networks and novel drug targets

**DOI:** 10.1101/2025.02.04.635308

**Authors:** Mylarappa Ningappa, Chethan Ashokkumar, Divya Murali, Dhivyaa Rajasundaram, Brandon W Higgs, William McDonald, Lori Schmitt, Liang Zhang, Ansuman Chattopadhyay, Matt Kane, Patrick Murphy, Kyle Soltys, George Mazariegos, Shakti Gupta, Upendra Kar, Timothy Billiar, Adriana Zeevi, Qingyong Xu, Miguel Reyes-Múgica, Shankar Subramaniam, Rakesh Sindhi

## Abstract

Acute T-cell-mediated transplant rejection, the earliest contributor to immunological graft failure responds variably to T-cell suppression. Early rejection can co-exist with donor-specific antibodies which also shorten graft survival, without affecting graft histology visibly. Here, during early liver transplant rejection, blood leukocytes manifest primed T-cells with heat shock protein (HSP) signaling and transcriptional programs for memory cell expansion. Corresponding biopsies demonstrate upregulated immune synapse and proteasomal genes in T-cell-infiltrated portal and central regions of the liver lobule. With donor-specific antibodies, the intervening intralobular region demonstrates a germinal-center-like allograft response with upregulated CD40, chemokine, T-follicular and complement signaling, which intensifies along the direction of sinusoidal blood flow from portal to central regions. Proteasomal and HSP90 inhibitors suppress donor-specific alloresponses of T- and B-cells in blood samples from patients with early rejection. The molecular injury spectrum of early rejection is complex, spatially differentiated and reveals novel early immunosuppressive strategies to extend graft survival.

## Introduction

Liver transplantation (LT) is life saving for children with congenital liver diseases and allows most recipients to achieve self-supporting adulthood^1^. The indications include cholestatic diseases in nearly three-fourths, metabolic diseases in a fifth, and unresectable liver malignancies in the remainder^2^. Nearly 80% of such recipients are transplanted before the age of ten years^3^. In contrast, the median age at LT during adulthood is 52 years^4^. However, a fifth of all pediatric LT recipients experience graft loss within 20-30-years after transplantation, the majority due to immunologic injury^5,6^. As a result, ∼40% of these recipients can require re-transplantation over the expected 80-year lifespan. Thus, lifelong survival of the transplanted liver in children presents a unique challenge and requires a fundamental advance in our understanding of allograft injury and the limited repertoire of immunosuppressants. New immunosuppressants have not been developed since approval of the costimulation blocker belatacept in 2011^7^.

The earliest immunologic injury to the allograft is T-cell-mediated rejection (TCMR)^8^. TCMR occurs in the first few weeks after LT despite use of Tacrolimus, an inhibitor of cytokine production by T-cells^9^. TCMR is characterized by lymphocyte-rich infiltrates around bile ducts or endothelial cells in the portal or central veins of the liver lobule. Up to half of all early TCMR events are accompanied by donor-specific anti-HLA antibodies, which do not alter graft histology further, unless antibody-mediated rejection (AMR) develops. Although deposition of the complement fragment C4d is used as a marker of AMR, it is not always reliable, and AMR remains underappreciated in liver grafts^10^. Because both TCMR and DSA predispose to late rejection and graft loss, the underlying mechanisms can define the molecular injury continuum connecting early and late allograft injury and reveal novel drug targets^8,11–13^. Graft histology offers no such mechanistic insights. As a result, anti-rejection drugs are used empirically risking cumulative overexposure, life-threatening infections, and post-transplant lymphoma^14^.

Here, we define immune-cell-based mechanisms of TCMR with single leukocyte whole transcriptome analysis (scRNAseq) of paired pre- and early post-transplant samples from children with LT. scRNAseq can uniquely ascribe mechanistic gene networks comprised of hundreds of genes to their origins in single immune cells^15^. Alternatives like single cell cytometry can assess far smaller numbers of protein markers. With bulk RNAseq, disease-associated gene expression originating from cellular mediators is diluted by non-specific expression from bystanders. Next, we define the spatial specificity of these mechanisms in the liver allograft using spatial transcriptomics (ST) of corresponding liver biopsies^16^. These findings are further validated with an independent biopsy cohort and the localization of key mechanisms confirmed with single cell spatial molecular imaging^17^.

## Results

The study design is shown in Figure 1.

**Figure 1.**
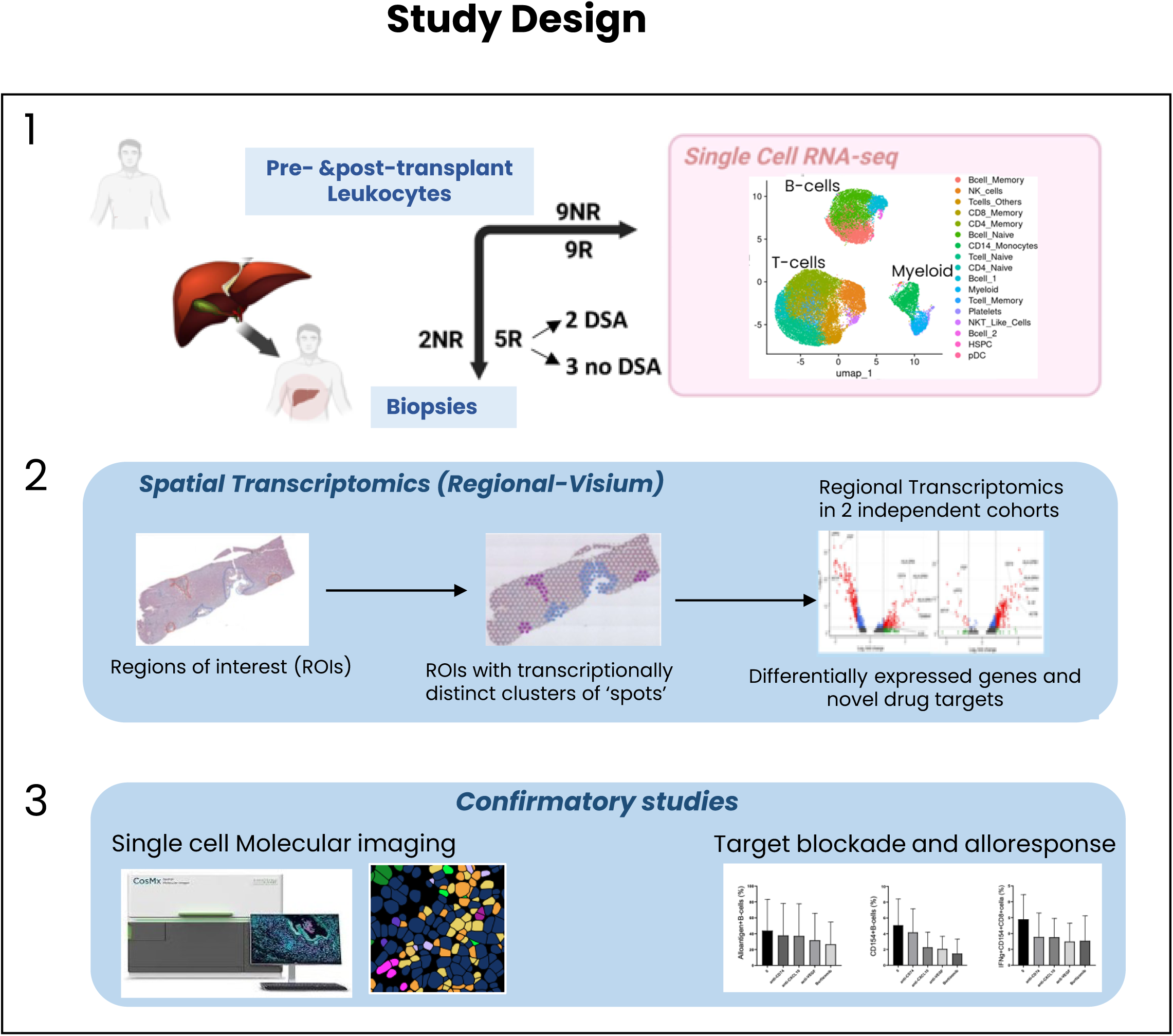
Study design. Single cell RNA sequencing was performed on leukocytes samples from 18 children, 9 with and 9 without rejection (R and NR). Spatial transcriptomics was performed on regions on corresponding biopsies from 2 Non Rejectors and 5 Rejectors of whom two also had circulating donor specific antibodies (DSA). Findings were confirmed with single cell spatial molecular imaging and functional immune assays.

### Memory cell priming and expansion

We obtained paired blood samples before and within the first 12 weeks after LT from eighteen children and annotated leukocyte subsets using reference datasets (Figure 1, Figure 2A, Table S1a-S1b) https://azimuth.hubmapconsortium.org/). Leukocyte subsets were compared between nine rejectors (R) with biopsy-proven ACR and nine non-rejectors (NR). Comparisons revealed higher counts of CD8 memory and naïve CD4-cells in pre-transplant samples and higher counts of memory subsets of B-cells, CD4 and CD8 cells in post-transplant samples from rejectors (Figure 2B). Several differentially expressed genes between rejectors and non-rejectors (DEGs, p-adj <0.05, Log2FC >0.25, Table S2-S3) were also identified in each leukocyte subset.

**Figure 2.**
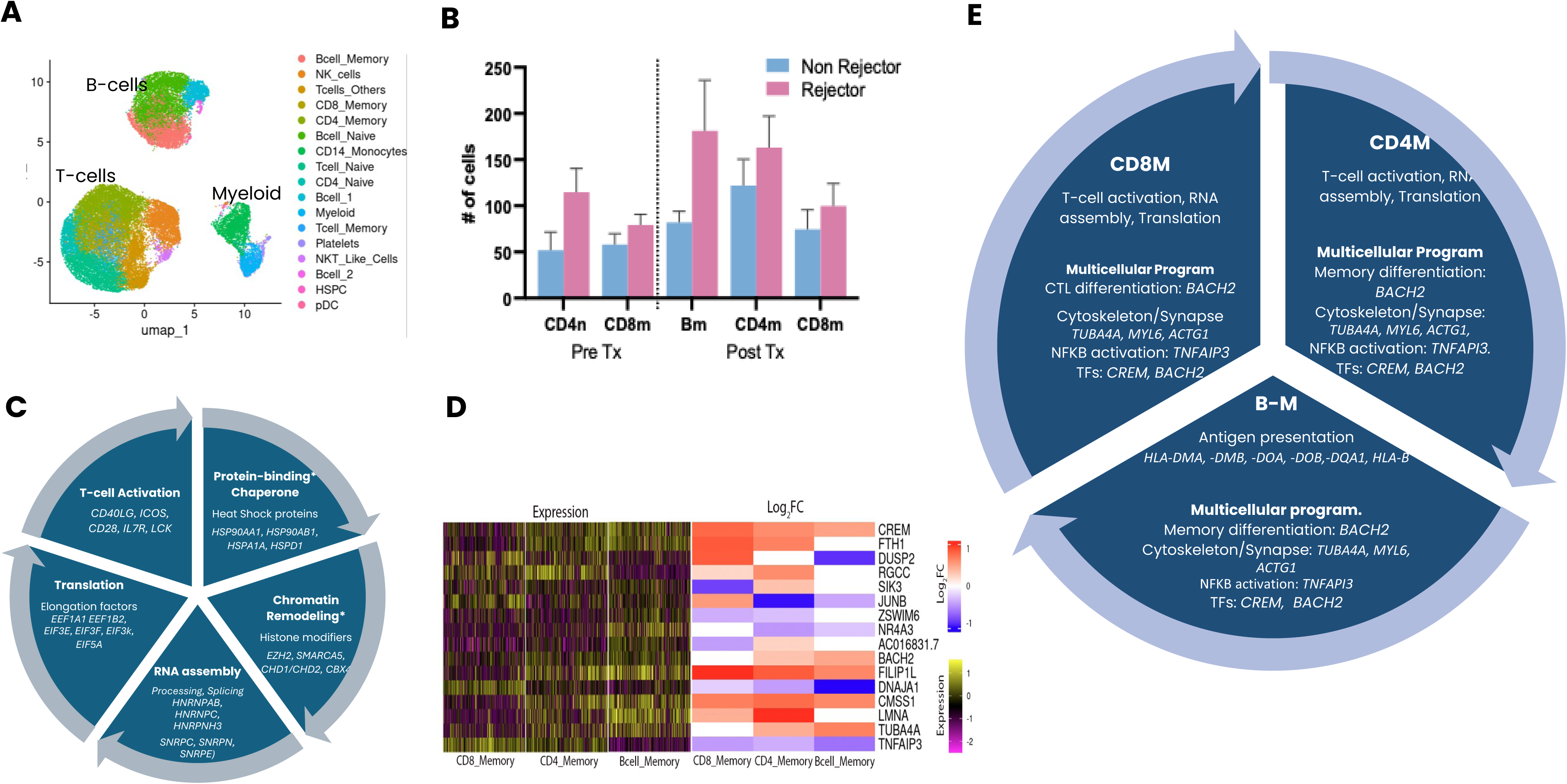
Leukocyte and allograft transcriptome of TCMR. **A**. UMAP shows leukocyte subsets in post-transplant samples. **B**. Counts of naïve (n) and memory (m) CD4, CD8 and B-cells in rejectors (R, pink bars) and non-rejectors (NR, blue bars) in pre-transplant and post-transplant leukocytes. **C**. Summary of upregulated functional gene sets in pre-transplant leukocytes. **D**. Heatmaps show covarying expression of genes in the M3 module of CD8 cells, in all three expanded memory cell subsets in all post-transplant samples (left) and differential expression of module genes in R compared with NR samples (right). **E**. Summary of multicellular transcriptional program genes underlying cell-specific function.

We first focused on expanded pre-transplant CD8 memory cells in rejectors. Pre-transplant donor-responsive CD8 memory cells which express CD40LG are known to predict clinical liver transplant rejection in an FDA-approved assay^18^. In support of primed memory T-cells, we found that upregulated DEGs in this subset clustered into functional categories involved in chromatin remodeling, transcription and translation, and T-cell activation (Figure 2C). These genes included the heat shock proteins (*HSP90AA1, HSP90AB1)*, which bind to and regulate folding of proteins like histones and chromatin-modifying genes *(EZH2, SMARCA5, CHD1/CHD2, CBX4)*; pre-mRNA processing (*HNRNPAB, HNRNPC)* and splicing *(SNRPC, SNRPN, SNRPE);* the eukaryotic translation elongation factors 1, 3 and 5 (*EEF1A1 EIF3E, EIF5A*): and T-cell activation (*CD40LG, ICOS, CD28)* (Figure S1, Table S2). The abovementioned genes for translation elongation, RNA splicing, and T-cell activation were also upregulated in the expanded naïve CD4 subset in pre-transplant samples from rejectors. After transplantation, T-cell activation genes remained upregulated in the expanded CD8 memory and CD4 memory subsets in rejectors (Table S3). In the expanded memory B-cells in post-transplant samples from rejectors, upregulated genes were noteworthy for those that drive antigen presentation via MHC class II (*HLA-DMA, -DMB, -DOA, -DOB)* and MHC class I (HLA-B) antigens (Table S2). Chromatin remodeling due to prior antigenic stimulation is known to prime T-cells to respond vigorously upon antigenic re-exposure^19,20^. HSP-induced priming and chromatin remodeling is also implicated in memory maintenance in other organisms^21^. HSP90 inhibitors suppress inflammation in plaque type psoriasis and could also prevent TCMR^22,23^.

### Multicellular transcriptional activation and memory differentiation in early TCMR

Among DEGs in the various subsets, we found that 172 genes in pre-transplant and 280 genes in post-transplant, were present in two or more leukocyte subsets (Figure S1D, Table S4). To determine whether this expression pattern represented multicellular transcriptional activation in TCMR after transplantation, we focused on identifying genes that covaried across the various leukocyte subsets with those in each of the expanded memory subsets in rejectors. These genes were organized into several modules based on covariance patterns in each cell type (Figure S2, Methods). For each module, we identified hub genes with the highest connectivity with others in that module (Table S5). Modules in which all genes were downregulated, and those containing ribosomal and mitochondrial genes were not considered.

TCMR involves coordinated responses of functionally distinct cell types, as illustrated by expression patterns of genes in module three of CD8 memory cells (CD8M3) in other memory cell populations (Figure 2D). Consistent with their post-transplant expansion, all three memory cell types show three upregulated genes: *CREM,* which promotes proliferation and activation of T- and B-cells^24,25^. *CMSS1*, a ribosomal protein, which enhances protein synthesis, and *FILIP1L*, which regulates cell proliferation^,26^. Further, *TNFAIP3* which suppresses NFKB-mediated inflammation is downregulated in all three memory cells^27^. The expression pattern of *BACH2* is consistent with its role in maintaining immunological memory when upregulated, as in memory CD4 and B-cells, and differentiation to effector CTL when downregulated, as in CD8 memory cells^28^. Upregulated *LMNA* and *FTH1* in CD4 and CD8 memory T cells are known to produce coordinated T-cell activation and infiltration at sites of inflammation, respectively^29,30^. Upregulation of *DUSP2*, which regulates cell proliferation has been reported previously in TCMR^31^. *CREM* is a known component of multi-cellular transcriptional programs in tumor-infiltrating immune effector cells^32^. The tubulin, myosin, and actin genes *TUBA4A, MYL6, ACTG1* in module BM1 (not shown) are respectively, components of the actin cytoskeleton which aids formation of the immune synapse. Thus, covarying expression of common genes in such functionally diverse leukocyte subsets can recruit associated inflammatory pathways in varying combinations explaining the varying histology and treatment response in individual patients with T-cell-mediated rejection (Figure 2E, Table S5).

### Portal and central signaling mechanisms in TCMR

TCMR is diagnosed if T-cell-rich infiltrates damage bile ducts in the portal regions of the liver lobule, with or without damage to adjacent portal venous or central venous endothelium^8^. To further determine injury networks in TCMR, we used Visium technology to examine ST of these lobular regions of formalin-fixed, paraffin embedded (FFPE) LT biopsies. Biopsies were obtained from a subset of test subjects, five with and two without rejection. Among 6 transcriptionally distinct clusters of ‘spots’, the cholangiocyte-gene-rich cluster 4 and the GLUL-gene-rich cluster 5 overlay portal and central liver lobular regions, respectively, as confirmed by the pathologist (Figure 3A, 3B, Figure S3, Table S6)^33^. These two regions contained 356 DEGs including 336 upregulated genes, and 552 DEGs including 157 upregulated genes, respectively, in rejection compared with non-rejection biopsies (Table S7-S8, Figure 3C, Figure S4). Downregulated genes were predominantly hepatocyte-derived metabolic genes, consistent with hepatocyte dysfunction during ACR. In the hepatocyte-rich central region, downregulated genes exceeded counts of upregulated genes which were mostly immune function genes. The largest functional clusters among these DEGs indicated a dominant IFNγ response (>21 DEGs) accompanied by leukocyte cell-cell adhesion (>58 DEGs), histone deacetylation (13 DEGs) or nucleosome dynamics (5 DEGs) in both regions, and mRNA processing (18 DEGs) and antigen presentation (7 DEGs) in the portal region (Figure S4A, B).

**Figure 3.**
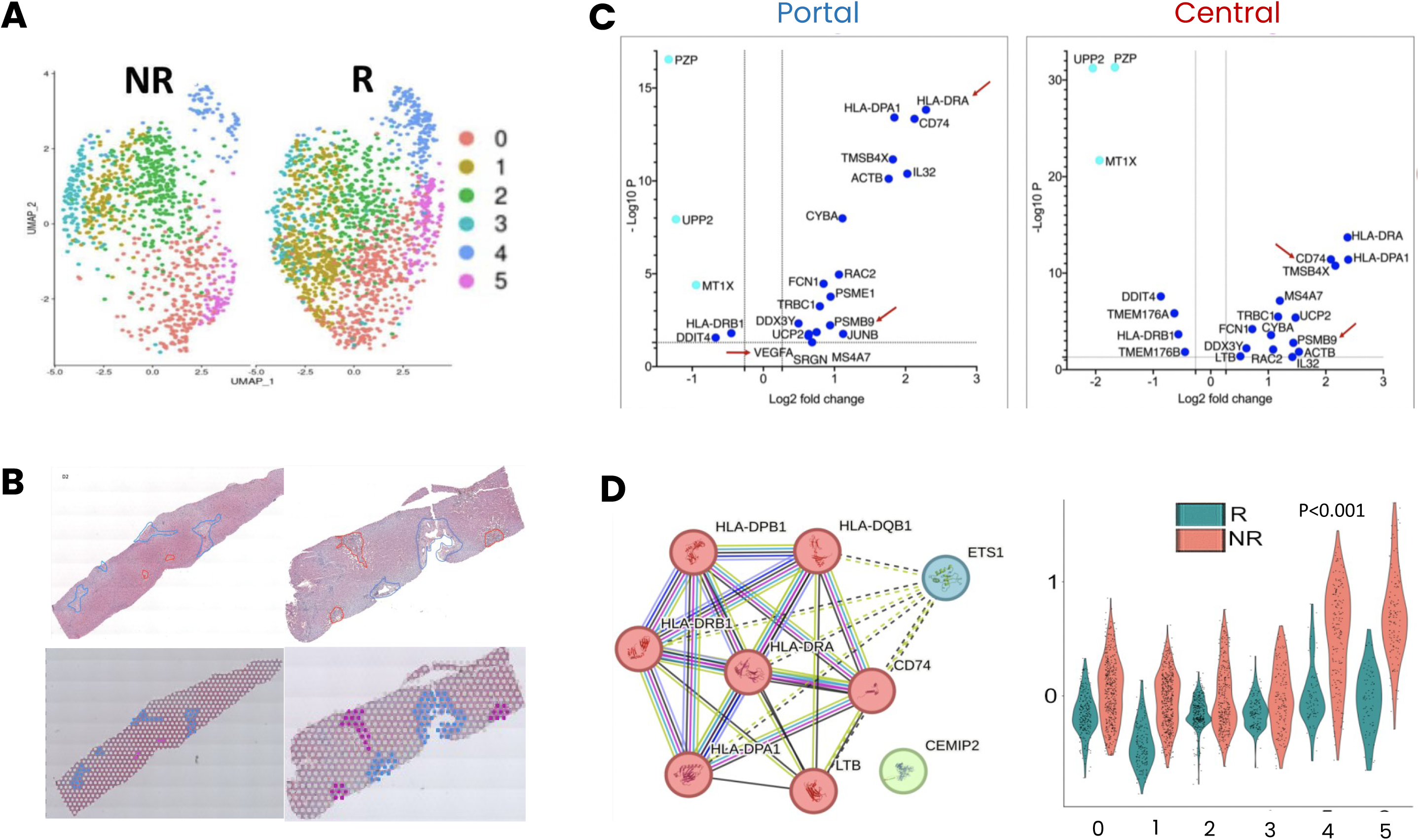
**A**. UMAPs show 6 clusters of ‘spots’ in an R and NR transplant biopsy. **B**. Hematoxylin-eosin-stained NR (upper left) and R biopsies (upper right) overlaid with Visium spots (lower panels, blue in portal and red in central liver lobular regions). **C**. Volcano plots show key differentially expressed genes including the drug targets, (red arrows) in portal (left) and central (right) regions of rejection biopsies. **D**. (left panel) A nine-gene network co-expressed with T-cells and dendritic cells. Right panel. Violin plots show expression of this 9-gene network in 6 clusters of spots (p-adj <0.001) in R (red violins) compared with NR (blue) biopsies. Clusters 4 and 5 overlie portal and central regions.

Among all DEGs, 146 genes in the portal and 96 genes in the central region were also differentially expressed in the various leukocyte subsets in post-transplant samples (Table S9). These common genes contained NFKB-regulated networks in both allograft regions (Figure S4C-D). Potential upregulated targets in these networks that could be blocked with known pre-clinical or clinical agents were, *CD74*, which aids antigen binding to MHC class II molecules, and the proteasome implicated by *PSMB9* in both regions (Figure S4C-D). *VEGF* was upregulated in the portal region only. The upregulated CD8A and the cytotoxin *GZMA* which characterize CTLs in both regions are accepted mediators of TCMR. Immune synapse formation was strongly suggested by the upregulated actin cytoskeleton gene *ACTB* and its stabilizer, *TMSB4X,* and a 9-gene network which included the recently described immune synapse genes, *CD74, HLA-DRA, LTB* in liver transplant biopsies^34^. These genes participate in MHC-dependent antigen presentation, were co-expressed with T-cells and classical dendritic cells in each biopsy and were differentially upregulated most in portal and central clusters in rejection compared with non-rejection biopsies (Figure 3D, Table S10). These findings were confirmed in seven independent liver transplant biopsies, six with and one without rejection (Table S11). Both regions of both rejection biopsy cohorts contained 31 upregulated DEGs including those for actin cytoskeleton (*ACTB, TMSB4X, FYB1, GBP1, LCP1)*, MHC II antigen-presentation (*HLA-DMB, CTSS, FCER1G)*, the cytotoxins/cytokine (*GZMA*, *GNLY, IL-32)*, and the proteasome (*PSMB9)* (Table S12). Memory T-cells were strongly implicated as well because the portal regions of both biopsy cohorts contained upregulated HSP genes including those found in pre-transplant CD8 memory cells in leukocytes from rejectors (*HSP90AA1, HSP90AB1*) (Table S13).

### Intralobular germinal-center-like allograft response during TCMR with DSA

Many early TCMR events are accompanied by cDSA, without deposition of the complement component C4d, which is seen in antibody-mediated rejection^10^. To determine whether cDSA was associated with molecular injury networks, we compared two of five rejection biopsies with cDSA, with the remaining three rejection biopsies without cDSA. UMAPs identified 6 clusters of spots in all rejection biopsies (Figure S5). Among them, clusters 2 and 4 located in intralobular regions showed differential expression of several humoral immune response genes (GO: 0006959) in rejection biopsies with cDSA compared with those without cDSA (Figure 4A and B, Tables S14, 15). These genes included the upregulated chemokines *CXCL10*, which primes CD8 cells to differentiate into CTL and promotes differentiation of activated B-cells into plasma cells^35,36^, and *CXCL13* which drives B-cells toward germinal centers and antibody formation^37^. Both chemokines can be targeted by blocking antibodies. We compared independent rejection biopsies, four with cDSA and two without cDSA by organizing spots into three regions, cholangiocyte-rich portal, GLUL-rich central, and the intervening intralobular region. We again found upregulated humoral immune response genes in the intralobular regions of rejection biopsies with cDSA including chemokines such as *CXCL10* (Table S16). Other upregulated genes were noteworthy for CD40 and NFKB signaling, and T-follicular cells, which characterize T-cell-mediated activation of B-cells in germinal centers (GC)^38–40^.

**Figure 4.**
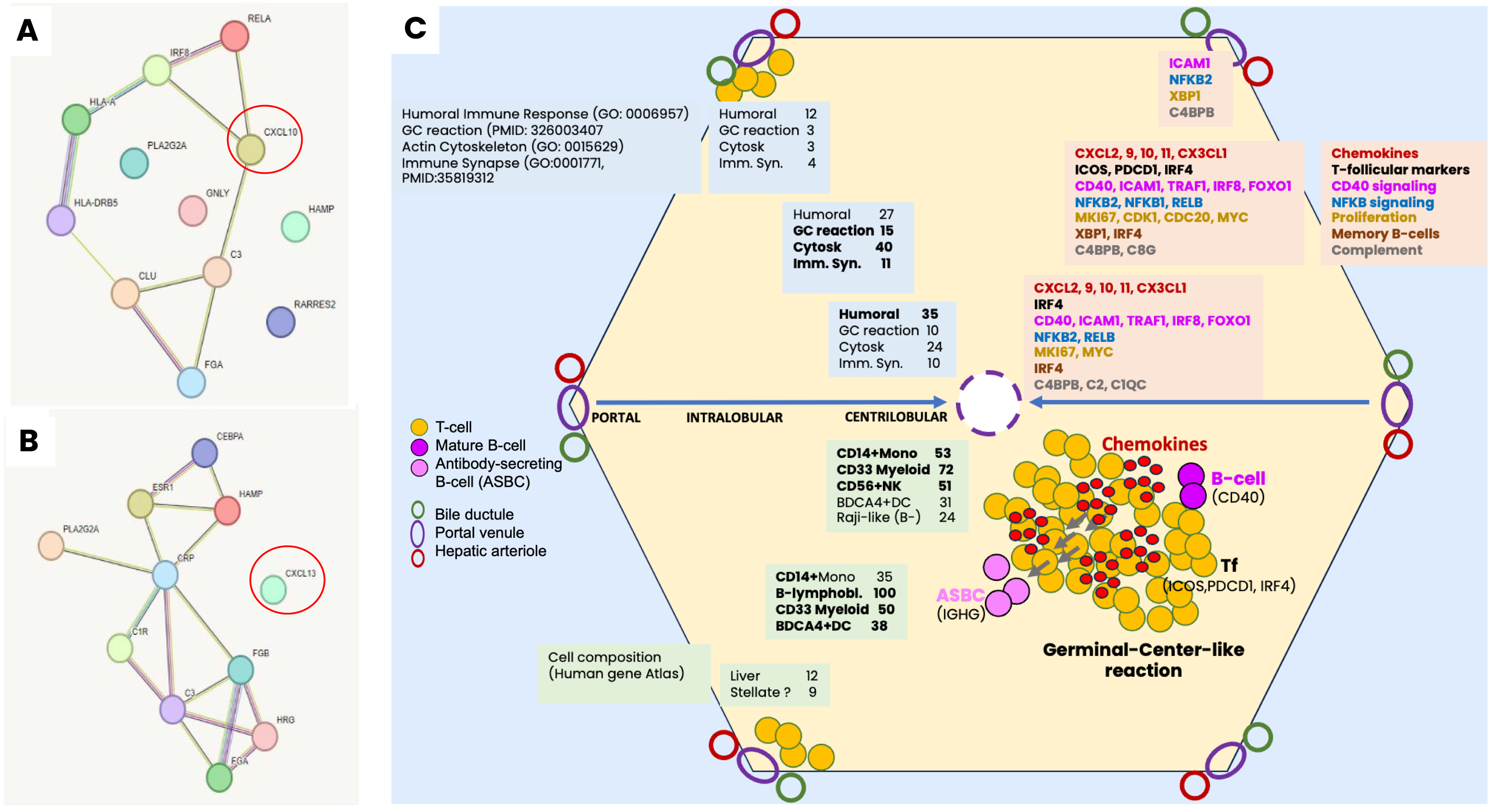
TCMR with cDS. **A. and B**. Differentially expressed humoral immune response genes (GO:0006959) in intralobular clusters 2 and 4 in rejection biopsies with cDSA. **C**. Schematic showing spatial organization of the germinal-center-like reaction in the liver lobule. Portal regions at corners contain bile ducts (dark green circle), portal venule (deep purple oval) and hepatic arteriole (red circle). Centrilobular region contains central vein (dashed purple outline). The intervening intralobular region contains liver sinusoids (blue arrow). Blue boxes list counts of differentially expressed genes (DEGs) in key pathways. Light orange boxes list selected pathway DEGs. Green boxes list counts of cell-specific DEGs. Key participants in the GC reaction are shown in the lower right quadrant of the hepatic lobule. ASBC=antibody-secreting B-cell.

Because these additional findings emerged in the larger numbers of rejection biopsies with DSA in the independent cohort, we maximized sample sizes for re-analyses by combining both cohorts. Gene counts for the GC-like reaction, immune synapse and cytoskeleton were lowest in portal regions, peaked intralobularly and declined to intermediate levels centrally (Figure 4C). The GC-like reaction genes included those for CD40 *(CD40, TRAF1, ICAM1)* and NFKB *(NFKB1, NFKB2, RELB)* signaling, proliferation (*MKI67, CDK1, CDC20, MYC*) and T-follicular (*ICOS, IRF4, PDCD1*) and memory B-cells (*XBP1, IRF4*)^38^. Together, these findings implicated an active intralobular GC-like reaction where chemokines promote cell migration and cell-cell interactions aided by the actin cytoskeleton and immune synapses. Alongside, we found that counts of humoral immune response genes doubled from 12 in portal to 27 intralobularly, and three-fold to 35 centrally, implying that the humoral injury network matures in the direction of sinusoidal blood flow from the portal region to the central vein. Among DEGs were cell-specific signatures (Human Gene Atlas) for CD33+myeloid cells which can include intrasinusoidal Kupffer cells and infiltrating CD14+monocytes, and the B-cell lineage in both regions. Gene signatures of CD56+NK cells, which are effectors of humoral injury were present centrally, consistent with a mature humoral injury network centrally.

To localize key cellular participants in TCMR with DSA, we performed a correlation of all unique DEGs across the expanded leukocyte subsets in post-transplant samples, with all genes expressed across a biopsy with TCMR and cDSA (Table S17-19). We found two gene modules (Figure 5A and B, Table S20). The CC1 module containing the T-cell genes *CD3D, CD3E, CD3G* localized to the portal and central regions, consistent with the diagnostic criteria for T-cell-mediated TCMR (Figure 5C). The CC2 module containing the B-cell genes, *CD79A* and *CD40* localized to the intralobular regions, coinciding with the location of differentially expressed chemokines and other humoral response genes (Figure 5D). This T-cell localization was also confirmed by correlations between DEGs in pre-transplant memory subsets of leukocytes and the same rejection biopsy. This relationship further confirmed the relevance of primed pre-transplant memory T-cells in facilitating rejection.

**Figure 5.**
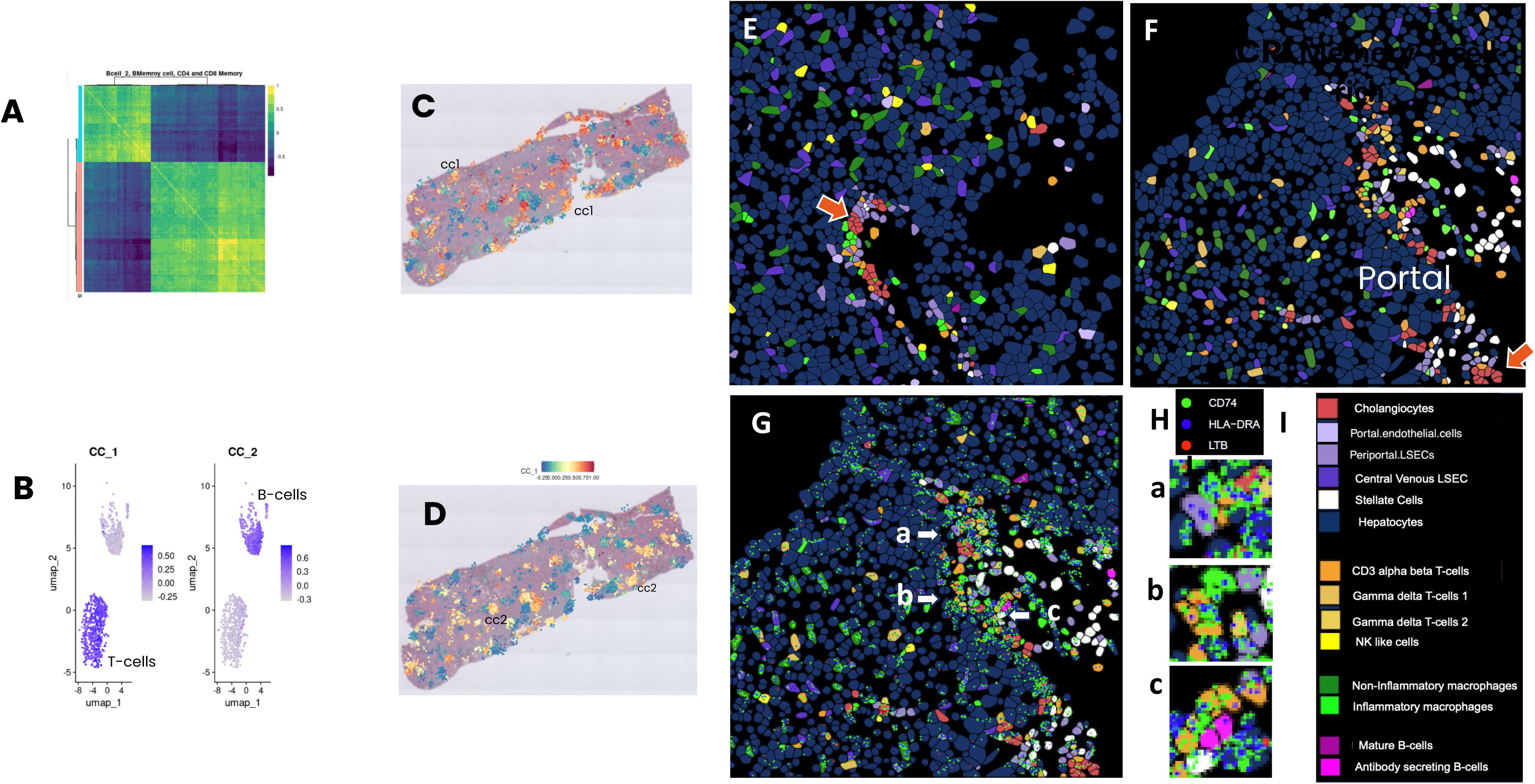
Cellular participants ‘in TCMR. **A**. Correlation of DEGs in post-transplant memory leukocyte subsets from rejectors and rejection biopsy with cDSA. **B**. UMAPs localizes modules cc1 and cc2 to leukocyte subsets. **C** and **D**. Localization of cc1 and cc2 modules in a rejection biopsy with cDSA. **E-I**. Spatial molecular images of 0.25 by 0.25 mm fields of view (FOV) with 1000-gene panel (CosMx). Portal regions containing cholangiocytes (orange arrows) in normal liver allograft pre-implantation (**E**), and liver allograft with transplant rejection (**F**), overlaid with colored dots representing immune synapse genes (**G**). **H**. Areas of cell-cell-contact, a-c in panel **G** are magnified to show immune synapse genes overlaid on T-cells in contact with **a**. portal endothelial cells, **b**. macrophage, and **c**. B-cells. **I**. Color legend for the different liver and immune cells.

### Confirmatory single cell spatial molecular imaging (SMI)

To localize injury pathways to single cells and regions of the liver lobule, we performed single cell SMI of FFPE liver sections from a pre-perfusion normal liver allograft and the liver allograft with rejection and cDSA using a 1000-gene panel^17^. Protein staining of cell membranes, epithelial cells and immune cells was performed with antibodies to B2M/CD298, pan-cytokeratins, CD68, and CD45, respectively. Single cells were annotated using reference scRNAseq datasets provided by Nanostring. In the portal and intralobular regions of the rejection biopsy with cDSA, we found the immune synapse genes *CD74, HLA-DRA, LTB* to be concentrated in regions of cell-cell contact between T-cells, and antigen-presenting cells like portal venous endothelial cells, macrophages and B-cells (Figure 5E-I). Also confirmed were higher frequencies of T-cells that expressed the memory markers *ITGAE, MK167, CD69,* and the cytotoxins *GZMA, GZMB*, *PRF1* (Figure S6). *The* intralobular region contained high-density expression of the chemokines *CXCL10, CX3CL1,* T-cells expressing T-follicular (Tf) cell markers, *ICOS, IRF4, PDCD1,* and B-cells which expressed *CD40*, and *IGHG2 (*Figure 6A-C). Interactions between Tf and germinal center B-cells via CD40-CD40L are essential for antibody production^41^. This inflammatory infiltrate was also accompanied by high-density expression of the complement gene *C1QC*, and minimal expression of the complement inhibitor, CD55 (Figure 6D). Interestingly, among 6 subjects with TCMR and cDSA, C1q binding to cDSA was strong in three, weak in two, with one showing no binding. Other GC participants expressed in this region included proliferation, (*MKI67, PCNA*) and memory B-cell genes (Figure S7). The directional maturation of the humoral immune response from portal to central regions, intralobular peaking of GC reaction genes and pathways, and the contribution of the macrophage-monocyte axis to TCMR with DSA observed with Visium transcriptomics and single cell SMI is summarized in Figure 4C.

**Figure 6.**
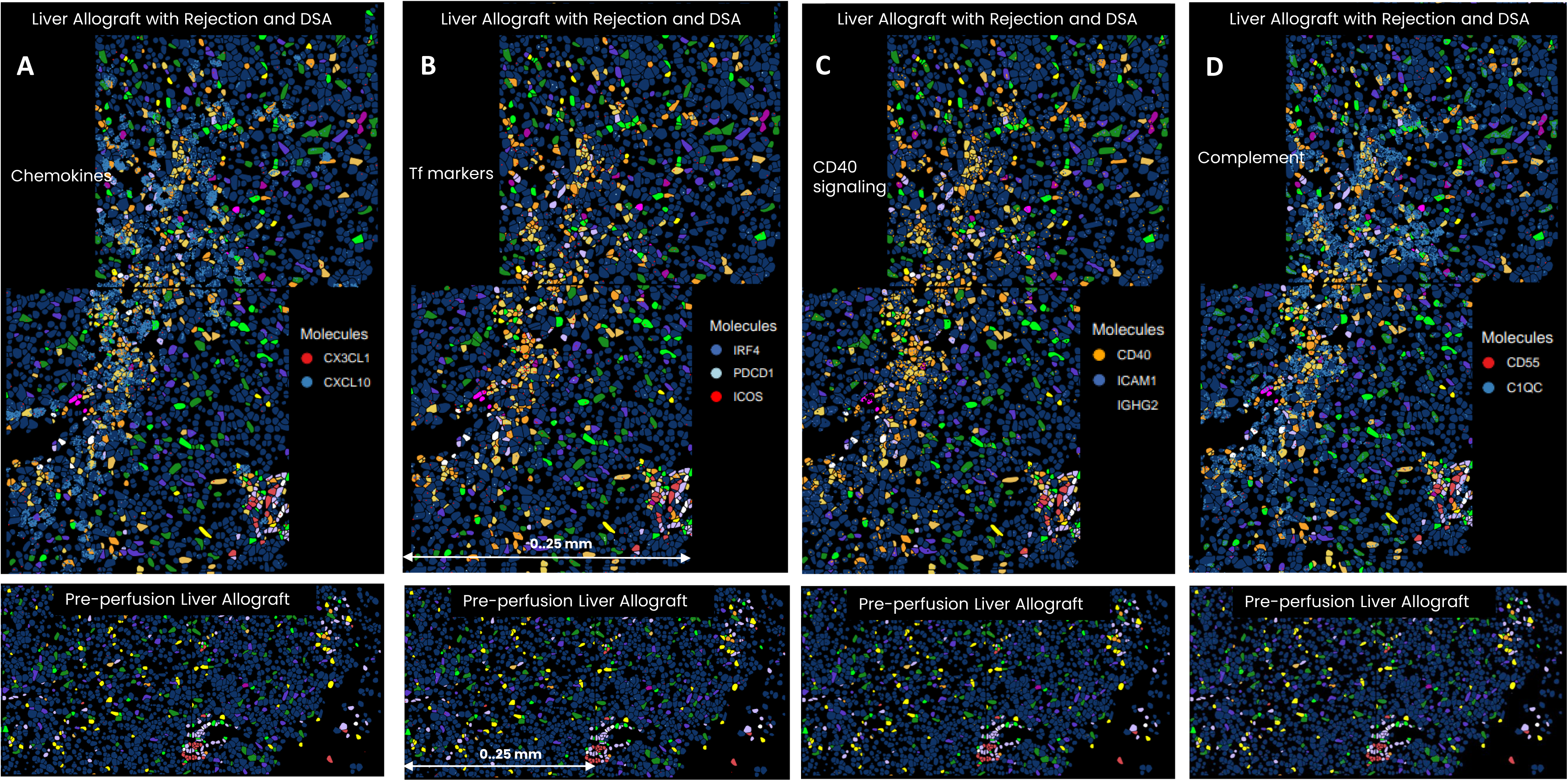
Germinal-center-like reaction in central and intralobular regions during TCMR with cDSA. (upper panels), **and pre-perfusion liver allograft** (lower panels). Components include from L to R, **A**) chemokines CXCL10 and CX3CL1, **B**) T-follicular cell markers (ICOS, PDCD1, IRF4), **C**) CD40 signaling (CD40, ICAM1) with expression of IGHG2 (black dots) in antibody secreting B-cells, and **D**) complement components C1QC and the complement inhibitor, CD55. Each image shows two FOVs, 0.025mm by 0.25 mm and was generated with the 1000-gene panel on the CosMx spatial molecular imager. Color legend for cells is shown in Figure 5. Color legend for genes is shown in boxes in each image.

### Target blockade suppresses the rejection alloresponse

We tested the effects of blocking the potential drug targets, CD74, CXCL10, VEGFA, and the proteasome on presentation of and effector response to HLA-mismatched alloantigen (Figure S8). These responses were measured with flow cytometry in co-cultures between three such pairs of HLA-mismatched responders and stimulators ^42–44^. Antibody-producing donor-specific B-cells are known to express IL-6 and CD40L (CD154) ^45,46^. CTLs co-express IFNγ and CD154^47,48^. Blockade of all targets except CD74 suppressed B-cell presentation of alloantigen and the B-cell and T-cell effector response (Figure S8). The proteasomal inhibitor, bortezomib suppressed all alloresponses consistently. To obtain preliminary evidence of potential clinical efficacy, we tested the in vitro effect of bortezomib on donor-specific alloresponses in leukocytes from three LT recipients experiencing rejection. Bortezomib suppressed presentation of donor antigen and donor-specific effector responses of B-cells and CD8 memory cells in leukocytes from these recipients (Figure 7A).

**Figure 7.**
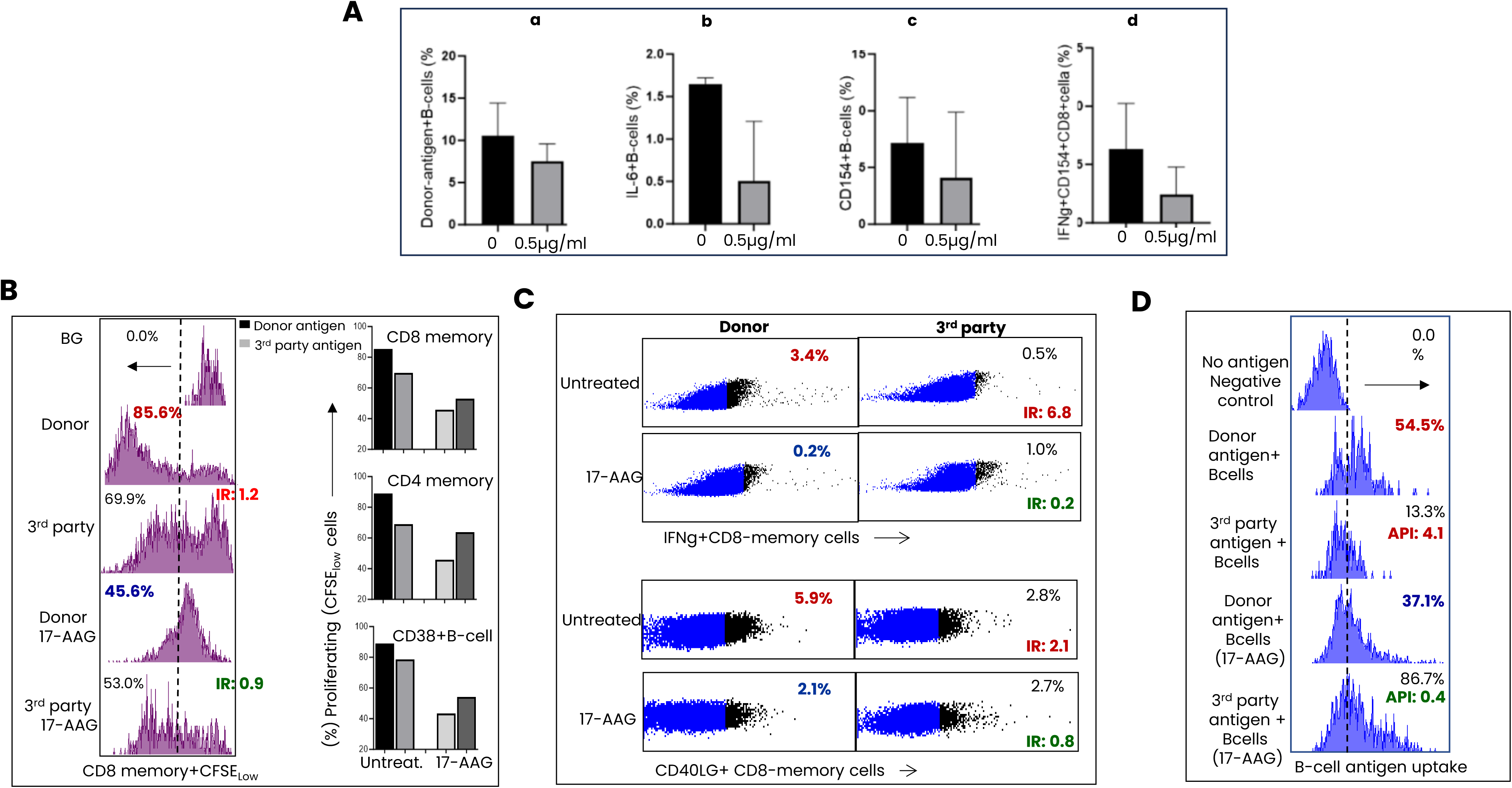
Effect of blocking novel drug targets on immune responses. **A**. Proteasomal inhibitor Bortezomib. Effect of proteasomal inhibition on the donor-specific immune response in leukocytes from three LT recipients experiencing TCMR. The immune responses are measured as **a**) the frequency of B-cells that present alloantigen, **b**) allospecific IL6+B-cells, **c**) allospecific CD154+B-cells, and **d**) allospecific IFNγ+CD154+CD8. **B**. 17-AAG suppresses donor-stimulated proliferation in TCMR. Left. Histograms show that 17-AAG inhibits proliferation measured as CFSE_low_CD8 memory cells to a greater extent after stimulation with donor cells (from 85.6% to 69.9%) than with third party (45.6% to 53%), The immunoreactivity index (IR) for CFSE^low^CD8 memory cells decreases from 1.2 to 0.9. Bar diagrams summarize these findings from the same co-culture for memory subsets of CD8, CD4, and B-cells (CD38+). **C**. 17-AAG suppresses donor-specific effectors CD8 memory cells in TCMR. Scatterplots show greater 17-AAG-mediated inhibition of donor-specific (upper plot) (%) IFNγ+ and (lower plot) (%) CD40LG+ CD8 memory cells, compared with third party-stimulated cells. the ratio of CD40LG+CD8memory cells induced with donor and third-party stimulation, termed the immunoreactivity index (IR) decreases from 2.1 to 0.8. **D**. 17-AAG suppresses B-cell presentation of donor antigen. Flow cytometry histograms show frequencies of B-cells from a recipient with TCMR, that present donor and third-party antigen, before and after treatment with 17-AAG. 17-AAG suppresses donor antigen presentation to a greater extent than third-party antigen. The ratio of these two measurements, or the antigen presenting index (API) decreases from 4.1 to 0.4.

*Preferential inhibition of donor specific alloresponses with the HSP90 inhibitor, 17-AAG:* We tested the effect of 17-AAG using modifications of the abovementioned donor specific alloresponses in leukocytes from LT recipients experiencing TCMR. Each alloresponse was measured after co-culture with leukocytes or antigenic lysate that was HLA-matched to donor (donor) or HLA-mismatched to donor and recipient (third-party). The results were expressed as the ratio of donor-induced and third-party-induced response and termed the immunoreactivity index (IR). For antigen presentation, the corresponding ratio was termed the antigen presenting index (API). Proliferation was measured with CFSE^low^ cells among memory (CD45RO+) subsets of CD4+ and CD8+T cells and memory (CD38+) B-cells. Pretreatment with 17-AAG inhibited proliferation and CTL responses to donor-stimulation (Figure 7B and 7C), and donor antigen presentation by B-cells (Figure 7D) to a greater extent than corresponding responses to third-party. The corresponding IR and API values which exceeded 1 before 17-AAG treatment changed to <1 after 17-AAG treatment.

## Discussion

Our studies of paired pre- and post-transplant blood leukocytes, and two independent biopsy cohorts show that early TCMR is defined by multicellular transcriptional activation of leukocyte subsets, and spatially differentiated injury mechanisms in allografts with TCMR and TCMR with cDSA. Reproducible mechanisms in biopsy cohorts are pre-requisites for clinical decision-making and a clear advance over allograft histology, or cytometry and bulk transcriptomics of leukocytes^49^. Primed CD8 memory cells expressing upregulated gene networks for chromatin remodeling such as HSP90 signaling, and T-cell activation are an important predisposing factor. After transplantation, memory CD8-, CD4- and B-cells expand, with multicellular activation of programs for differentiation of memory and cytotoxic T-cells, MHC-dependent antigen presentation, and the actin cytoskeleton. Consequences include cell migration, cell-cell interactions such as the immune synapse, and cytotoxic injury to the allograft. The blood leukocyte transcriptome is replicated in and reorganizes into spatially differentiated injury networks in the allograft. During TCMR, the T-cell-rich portal and central regions shows enhanced proteasomal signaling, immune synapse formation and CTLs. In our study, this localization has been guided by the well-established histology of TCMR^8^.

In TCMR with cDSA which is not accompanied by complement deposition, additional histological changes are not established. Here, localization of associated inflammatory networks has been aided by consolidating the lobular regions into cholangiocyte-rich portal, GLUL-rich central, and the intervening intralobular region. In this variant of TCMR, upregulated genes strongly implicate an active GC-like reaction in the intralobular and central regions. Participants include genes for several chemokines (*CXCL10, CXCL2, CXCL9, CXCL11 and CX3CL1*), T-follicular cells, CD40 and NFKB signaling and proliferation. The numbers of genes for the GC-like reaction and the humoral immune response increase from the portal to the central regions, in the direction of sinusoidal blood flow. The associated increase in cell-specific genes is consistent with the increasing engagement of the macrophage-monocyte lineage of cells such as Kupffer cells which line the sinusoid, with cells of the B-cell lineage, and T-cell-rich infiltrates. Chemokine production, chemokine-induced immune cell migration into the lobule are plausible outcomes of this engagement which can foster smoldering injury if left unchecked. Single-cell spatial imaging of a biopsy with TCMR and cDSA confirms this localization of cells and genes for the GC-like reaction, which facilitates antibody production. A final targeted immunosuppression strategy is the possible use of C1q esterase inhibitors in TCMR with cDSA and high-density expression of the complement component C1q centrally. This finding can explain smoldering antibody-mediated injury despite normal allograft function due to C1q-binding cDSA in patients with TCMR. Chemokines, GC-like reaction genes, and C1q can also serve as biomarkers for this TCMR variant. We are conducting spatial molecular imaging of additional biopsies to determine whether TCMR with cDSA has overlapping features with and thus reflects a path to frank ABMR.

An exciting finding in our study is the potential to use the proteasomal inhibitor bortezomib as an alternative to high-dose steroids to treat early TCMR after liver transplantation. Proteasomal inhibitors have been well tolerated when treating ABMR but have not been used for TCMR. In vitro suppression of donor-specific leukocytes from patients experiencing TCMR with bortezomib strongly supports follow-up clinical investigation as an alternative to high-dose steroid treatment of TCMR, which can cause hypertension, hyperglycemia, and weight gain. Among other potentially novel immunosuppressants, inhibition of HSP90 and related signaling mechanisms present a promising avenue to develop specific anti-rejection drugs. These agents suppress the donor-specific alloresponse preferentially attesting to the fundamental role of memory cells in TCMR and may be especially suited for sensitized recipients as alternatives to current lymphocyte depleting strategies. the germinal-center-like reaction in rejection with cDSA. Thus, our report offers the potential to fundamentally advance the understanding and early management of liver transplant rejection with targeted drugs which can extend graft survival.

## Materials and Methods

### Human Subjects

Study procedures were performed under University of Pittsburgh Institutional Review Board approved study #19030279 and #19050089. Subjects included 11 males and 7 females, mean ± SEM age 10.97 ± 2.57 years, of whom 9 experienced biopsy-proven early ACR based on Banff criteria (rejectors, R), and 9 who did not experience ACR (non-rejectors, NR). The age and gender distribution between groups was not significantly different (p=NS, Fisher’s exact test). Rejection was graded as mild in eight and moderate in one^10^. Blood samples were collected 15.66±4.76 days (mean ± SEM) before the rejection event in R. The indications for LT were biliary atresia-5, maple syrup urine disease-7, cystic fibrosis-1, cirrhosis due to total parenteral nutrition-1, propionic acidemia-1, primary sclerosing cholangitis-1, Alagille’s syndrome-1 and autoimmune liver disease-1. In the 90-day period after LT, anti-HLA donor-specific antibody (DSA) were identified in 5 of 9 R and 1 of 9 NR (p=0.04, Fisher’s exact test). The mean±SD fluorescence intensity of DSA measured with single antigen coated beads was higher among R, 5100±1591 and occurred earlier after LT, 34±4 days compared with MFI 3090, 75 days after LT in one NR. DSA resolved within 12 months after LT. DSA were characterized with single antigen coated beads (Luminex)^51^.

In the Viscum validation cohort, biopsies from nine subjects including 6 males and 3 females were included of which 7 were R and 2 were NR.,. The mean age ± SEM was 6.95 ± 2.11 years. The age and gender distribution between groups was not different(p=NS) Rejection was graded as mild in five and moderate in two^51^. The biopsy samples were collected at the time of rejection event in R. The indications for LT were hepatoblastoma-2, biliary atresia-1, hepatopulmonary syndrome (primary biliary atresia)-1, maple syrup urine disease-1, progressive familial intrahepatic type 2 cholestasis-1, propionic acidemia-1, methyl malonic acidemia-1 and staged palliation to extracardiac fontan-1.

Within 90 days post-LT, DSA were detected in 5 out of 7 R and 1 out of 2 NR (p=0.99, NS, Fisher’s exact test). The peak DSA fluorescence intensity, measured using single antigen-coated beads, was 9210 ± 5774 (mean ± SD) in R, occurring at a mean of 29 ± 9 days after LT, compared to an MFI of 22,500 at 55 days post-LT in the one 1 NR. Two of nine biopsies were excluded for failure to meet quality thresholds.

## METHOD DETAILS

### Sample collection

PBMC were obtained from whole blood collected in sodium heparin tubes, before and within 90 days after LT from all 18 children, Spatial transcriptomics (ST) was performed in corresponding formalin-fixed paraffin-embedded (FFPE) biopsies from 5 of 9 R and 2 of 9 NR (Viscum, 10XGenomics). In the Viscum validation cohort, spatial transcriptomics was performed in FFPE biopsies from 7 R and 2 NR within 90 days after LT. All biopsies were performed for suspected rejection based on elevated liver function tests and absence of vascular occlusion or bile duct obstruction on imaging studies. Confirmatory spatial transcriptomics was also performed on eight independent biopsies, 6 with rejection and 2 without rejection. For confirmatory single cell spatial molecular imaging (SMI) (CosMx, Nanostring), we selected one biopsy with ACR and cDSA from the first cohort, and a pre-perfusion liver allograft biopsy from a normal healthy organ donor.

### Single Cell RNA sequencing (scRNA)

Single cell RNA sequencing (scRNA-seq) libraries were generated for single cell RNA sequencing using Chromium 3’ v3 scRNA-seq Library Kit (10X genomics, CA) according to the manufacturer instructions at the Single Cell Core lab at the University of Pittsburgh. 4500-5000 PBMC cells were used per sample, and data yield approximated 2000-2500 cells/sample. 12 samples were multiplexed per pool. The libraries were sequenced in the UPMC genome Center using Novaseq 6000 instrument (Illumina, CA) to get up to 20,000 reads per cell.

### Spatial transcriptomics of (ST) FFPE biopsies using Visium platform

Sample preparation was performed with the Visium platform with the FFPE Gene Expression Starter Kit, Human Transcriptome (10x genomics, CA). FFPE sectioned tissue was placed on a 6.5mm x 6.5mm capture area of the 10XGenomics Visium slide with 5,000 barcoded spots, each containing millions of oligos include a 16 nucleotide (nt) spatial barcode for location, a 30 nt poly(dT) sequence for mRNA capture, a 12 nt Unique Molecular Index (UMI) unique for each transcript bound by the probe, and an Illumina Read 1 sequence for use in library preparation and sequencing. Tissue was deparaffinized and stained with hematoxylin-eosin. Microscopic imaging was performed and subsequently used to overlay gene expression patterns. Stained tissue was de-crosslinked to release RNA, specific probes covering the whole transcriptome were added to capture free mRNA targets, and probe pairs that have hybridized to RNA were ligated using a ligase. Tissue was treated with RNase and permeabilized to release ligation products which were captured on the Visium slides. Captured ligation products were extended by a polymerase to incorporate a complement of the Spatial Barcode sequence and UMI. The spatially barcoded, ligated probe products were released from the slide and PCR-amplified using common sample-indexing primers. The final library was sequenced at a depth of 25k raw reads per capture spot covered by tissue in most FFPE samples in the UPMC genome Center using Novaseq 6000 instrument. Sequencing data was aligned with the image using barcode information for location using SpaceRanger, (10x Genomics, CA). Each barcoded spot was 55um in diameter with 100um center to center distance between spots. Thus, there was an average of 5-10 cells per spot depending on tissue cellular density. Images were visualized with Loupe Browser.

### Spatial Molecular Imaging (SMI) of FFPE biopsies using CosMx

The FFPE samples were processed using the Human Universal Cell Characterization 1000-plex panel following the manufacturer’s protocol (Item number:121500005, NanoString Technologies, Seattle, WA), Tissue sections were cut to a thickness of 5 μm and mounted on Superfrost Plus Micro Slides within a 15 mm x 20 mm imaging area, and baked overnight at 60°C to enhance adherence. After deparaffinization, target retrieval was performed at 95-100°C for 15 minutes, followed by permeabilization with 3 μg/mL Proteinase K at 40°C for 30 minutes. Fiducial markers were applied for precise image alignment, followed by post-fixation with neutral buffered formalin and blocking with N-hydroxy succinimide (NHS)-acetate. The slides were then hybridized overnight at 37°C with RNA-specific probes from the 1000 plex RNA panel. Following stringent washes to remove unbound probes, DAPI staining was used for nuclear visualization, and cell segmentation markers CD298/Beta-2-Microglobulin (B2M), additional cellular markers, CD68, PanCK, and CD45 were applied. The prepared slides were loaded into the CosMx SMI instrument (Seattle, WA) for imaging. During imaging, branched fluorescent probes were hybridized to amplify signals, enabling the detection of 1,000 RNA targets within individual cells. The raw images were processed and decoded using the AtoMx™ Spatial Informatics Platform (SIP), a cloud-based service that provided detailed analysis and visualization. This allowed for comprehensive insights into cellular RNA expression profiles and spatial distribution, essential for understanding tissue architecture and cellular interactions.

### Statistical Analysis for scRNA-seq

All statistical analysis was performed in R version 4.2.1^52^. Cell Ranger (10x Genomics Cell Ranger version 7.2) was used to demultiplex, align (hg38), and quantify reads from the PBMC scRNA-seq samples. Filtered matrices were converted into SeuratObjects and downstream analysis was performed using Seurat (v.3.2.2).^53^ Low quality cells were filtered such that 1000 < nFeature_RNA > 150, and mitochondrial percent < 20. Each sample was normalized using SCTransformation. All samples were integrated using CCA integration with nfeatures = 3000. Dimensionality reduction was performed using PCA and visualized using UMAPs. Integrated Seurat object was projected to publicly available Azimuth’s Human PBMC reference^54–58^ Finally, the Seurat clusters were annotated with maximum cell type population. Differentially expressed genes (DEGs) for each cluster were identified for R versus NR in both Pre and Post LT conditions respectively (|LFC| >0.25, adjusted p-value<0.05). DEGs was also obtained in R for each cluster comparing DSA versus No-DSA in Post and Post LT condition.

### Functional Enrichment

We performed biological functional enrichment analysis on the differentially regulated genes within each cell population using multiple enrichment tools and algorithms, i.e.,), enrichR and further refined with protein-protein interactions in STRING.^59^; and gene set enrichment analysis using clusterProfiler 4.0^60,61^. ChIP-seq data from ENCODE^62^ was evaluated to confirm TF binding motifs within 3KB of the transcription start site. We searched for predicted upstream regulators of differential expression of genes (DEGs) in each cell cluster, used the Ingenuity Pathway Analysis “Upstream Analysis” tool (Kramer), and selected “Transcriptional Regulator” as a “Molecule Type” filter term to identify statistically significant upstream regulatory Transcription Factors (TF)^63^. Statistical significance was calculated using the right-tailed Fisher Exact Probability Tests; TF showing a p-value < 0.05 were considered statistically significant. The activity status of TF was determined by calculating the activity z-score, which is a statistical measure of how closely the expression pattern of TF-target genes present in the uploaded dataset compares to the expected pattern based on the literature findings. A positive score indicates an overall increase in activity of the TF and vice versa.

### WGCNA Analysis

We performed weighted gene co-expression network analysis to identify the common transcriptional programs in Pre and Post LT PBMC data^64^. Co-expressed gene modules were generated using hdWGCNA (High-Dimensional Weighted Gene Correlation Network Analysis)^65^. Cell types that showed significant increase in the number of cells in R compared NR were chosen to perform this analysis. Differentially expressed modules (DMEs) were identified between R and NR for downstream analysis.

We investigated the spatial-relevant gene coexpression modules using the SCoexp module from CellTrek^66^. For cells of interest, we first calculated a spatial-weighted gene cross-correlation matrix using the cell-gene expression matrix. Co-expression module detection was done using the consensus clustering approach with a default K through 2 to 8 and repetition of 20. For the identified co-expression modules, we calculated the cell-level module activity score and then investigated the module scores at the spatial level. Using cellTrek we mapped the resident immune cells of the liver and found modules within cells of interest.

### Immunostaining

To further identify molecular and cellular interactions, we performed immunofluorescence staining of CD74 and CD68 markers to confirm localization indicated by DEGs and co-expression analysis of Visium data.

### Flow cytometry assays

Whole blood or frozen Peripheral blood leukocytes (PBL) were incubated alone (No drug), with anti-CXCL10 at 37.7ug/ml, with anti-VEGF at 22.5ug/ml, with Bortezomib at 0.5ug/ml, concentration and incubated overnight at 37°C. After incubation, whole blood was ficolled and PBL’s were isolated. The PBL’s were divided into three aliquots after incubation and were used to study B-cell antigen presentation, and for lymphocyte co-culture assays to measure effector function of T- and B-cells.

#### Antigen presentation

PBL were stimulated with dye-labelled antigen as described earlier.^42,43^. Briefly PBMC were incubated with dye-labeled antigen for one hour at 37°C with 5% CO2. After incubation, the cells were labelled with anti-CD19 antibody to identify B-cells and acquired on flowcytometry. 7AAD was added before acquisition to separate live cells from dead cells. Flowcytometry data were analyzed using the gating strategy described earlier^43^.

#### Lymphocyte co-culture assay for effector T-cells

Peripheral blood leukocytes (PBL) were incubated with HLA mismatched stimulators and incubated overnight as described^18^. Briefly Responder and Stimulator PBL’s were surface labelled with anti-CD8APCH7 and anti-CD8PeCy7 respectively to distinguish responder T-cytotoxic cells from Stimulator T-cytotoxic cells. After incubation, cells were labelled with anti-CD3Percp5.5 to label T-cells. The cells were subsequently fixed and permeabilized using an Intracellular staining buffer set (Cytofix / Cytoperm kit; BD Biosciences), stained with anti-CD154BV421 (BD biosciences) and anti-IFNγ FITC (BD biosciences) and analyzed by BDBiosciences FACS CANTO II flow cytometry. Flowcytometry data were analyzed using the gating strategy described earlier^18^.

#### Lymphocyte co-culture assay-B-cells

Responder and Stimulator PBL’s were surface labelled with anti-CD19APCH7 and anti-CD19PeCy7 respectively to distinguish responder B-cells from HLA mismatched Stimulator B-cells and co-cultured together with 1:1 ratio and incubated overnight at 37°C and at 5% CO2. After incubation cells were labelled with anti-CD3Percp5.5 to label T-cells and CD16/56Percp5.5 to label NK/NKT-cells. The cells were subsequently fixed and permeabilized using an Intracellular staining buffer set (Cytofix / Cytoperm kit; BD Biosciences), stained with anti-CD154BV421 (BD biosciences) and anti-IL6PE (BD biosciences) and analyzed by BDBiosciences FACS CANTO II flow cytometry. Flowcytometry data were analyzed using the gating strategy described in Figure S9.

#### Lymphocyte co-culture assay to measure proliferative alloresponse

Responder PBL’s were incubated with HLA matched (donor) and HLA mismatched (third-party) stimulators and incubated overnight as described earlier^-67^. Responder PBL’s were labelled with Carboxyfluorescein succinimidyl ester (CFSE) dye. Stimulator donor and third-party stimulator leukocytes were irradiated. Cells were incubated at 1:2 ratio (Responder: Stimulator) for 5 days at 37°C with 5% CO2. After incubation cells were surface stained with anti-CD3 PE, anti-CD8-PeCy7, anti-CD45RO-PeCy5, anti-CD19-APCH7, and anti-CD38-APC and analyzed by BDBiosciences FACS CANTO II flow cytometer using the gating strategy described earlier^67^. All antibodies for flow cytometry were obtained from BDBiosciences (CA). CFSE dye was from Invitrogen, Thermo Fisher (MA).

### Statistical Analysis Spatial Transcriptomics: Data processing

Gene count matrices from the Space Ranger pipeline were converted into a Seurat object and downstream analysis was performed using the R package, Seurat (v.3.2.2)^44^. Pre-processing of the ST datasets include removing of spots with more than 8% mitochondrial genes or fewer than 150 detected gene counts. The SCTransform normalization was used to normalize the gene-spot matrices followed by screening of the hypervariable genes. The objects from each of the biopsy sample were integrated using the Seurat anchor-based integration workflow followed by dimensionality reduction on 2000 variable genes. The identified clusters were visualized in low-dimensional space using Uniform Manifold Approximation and Projection (UMAP), and the corresponding spot coordinates were visualized by superimposing on the images. Differentially expressed genes (DEGs) between the R and NR was identified, for each cluster with a threshold (adjusted P-value < 0.05, |log2 FC| > 0.25) based on Benjamini-Hochberg correction.

### Cell-type deconvolution analysis

Publicly available Human scRNA-seq liver cell atlas data (accessed 2022-04-01)^17^ was used as reference data for deconvolution of Visium slides. Thereafter, we implemented Conditional Autoregressive Model-Based Deconvolution (CARD)^68^ for spatially informed cell type deconvolution based on cell type specific expression using the reference scRNA-seq data. Briefly, the CARD object was created using the sparse matrix of the raw scRNA-seq count data, a data frame containing the single cell metadata information, sparse matrix of the raw spatial resolved transcriptomics count data, and the spatial location with the x and y coordinates. The cell type proportion matrix was jointly visualized as the scatter pie plot. We then facilitated the construction of a refined spatial tissue map, and the cell type proportion of individual markers genes were visualized at an enhanced resolution.

### Approach to analyzing regional transcriptomes

*W*e localized potential regions of interest (ROIs) by: *a)* identifying distinct cell clusters and genes unique to each cluster, and b) evaluating each cluster for cholangiocyte-specific and central-region-specific genes described previously^69^. The two ROIs where the lymphocyte-rich infiltrate of TCMR include the cholangiocyte-rich portal region, and the pericentral region with high expression of the glutamate-ammonia ligase gene, *GLUL*.

### CosMx Single cell spatial transcriptomics analysis

A total of 30 fields of view (FOV) were captured from the sample within the imaging area for the 1000-plex panel. Segmentation was performed using cellpose^70^ with default parameters. Based on the segmentation results, 343,081 cells were identified. cell boundaries obtained helped assign transcripts at the cell-level and obtain a transcript by cell count matrix^68^. Cells with an average negative control count greater than 0.5 and less than 20 detected features were filtered out. QC was performed at the level of FOVs and cells. FOVs that had low performance including abnormal segmentation, blurry FOVs, <80 counts/cell were removed. Cell level QC was performed by removing low count cells (<20 counts), doublets and multiplets (area > mean± 5 SD) Cells were normalized using total count. Dimensionality reduction was performed using PCA (npcs =50). We obtained 18 clusters using Leiden cluster with resolution 0.2. neighbor network was performed with Jaccard cutoff: 0.067. Annotation was performed by taking consensus between insitu type using semi supervised approach^68^ and Azimuth and Liver reference datasets ^54–58^. To identify subpopulation markers, we ran findMarkers (scran package, version 1.22.1)^71^, blocking by sample, considering highly variable genes only, and testing for positive log-fold changes.

## Acknowledgements

Funding: Raizman Haney Endowed Fund-RS, Hillman Foundation of Pittsburgh-RS, Marjory K. Harmer endowment for research in Pediatric Pathology (MRM)

## Author contributions

All authors reviewed the final datasets and contributed to writing the manuscript. RS obtained funding, conceived, and directed the project. SS supervised all analyses. MN coordinated the project and performed CosMx ST and scRNAseq, CA performed Flowcytometry and scRNAseq, WM performed Visium ST, BWH and SG conducted scRNAseq analysis, DR and DM conducted Visium and CosMx analysis. LZ and MK reviewed CosMx analysis, PM reviewed Visium analysis, and AZ and QX carried out DSA analysis. MR integrated biopsy and Visium analysis. and QX and AZ generated and analyzed donor-specific antibody data, LS prepared ST samples. AC performed network analysis. UK and TB reviewed the data.

## Data availability

RNA-seq data (raw and processed) and CosMx data (flat files) will be deposited in the public database ‘’Gene Expression Omnibus (GEO)’’. GEO access will be released once the manuscript has been accepted for publication.

## Supplementary Materials

**Legends for Supplementary Figures**

**Figure S1.**
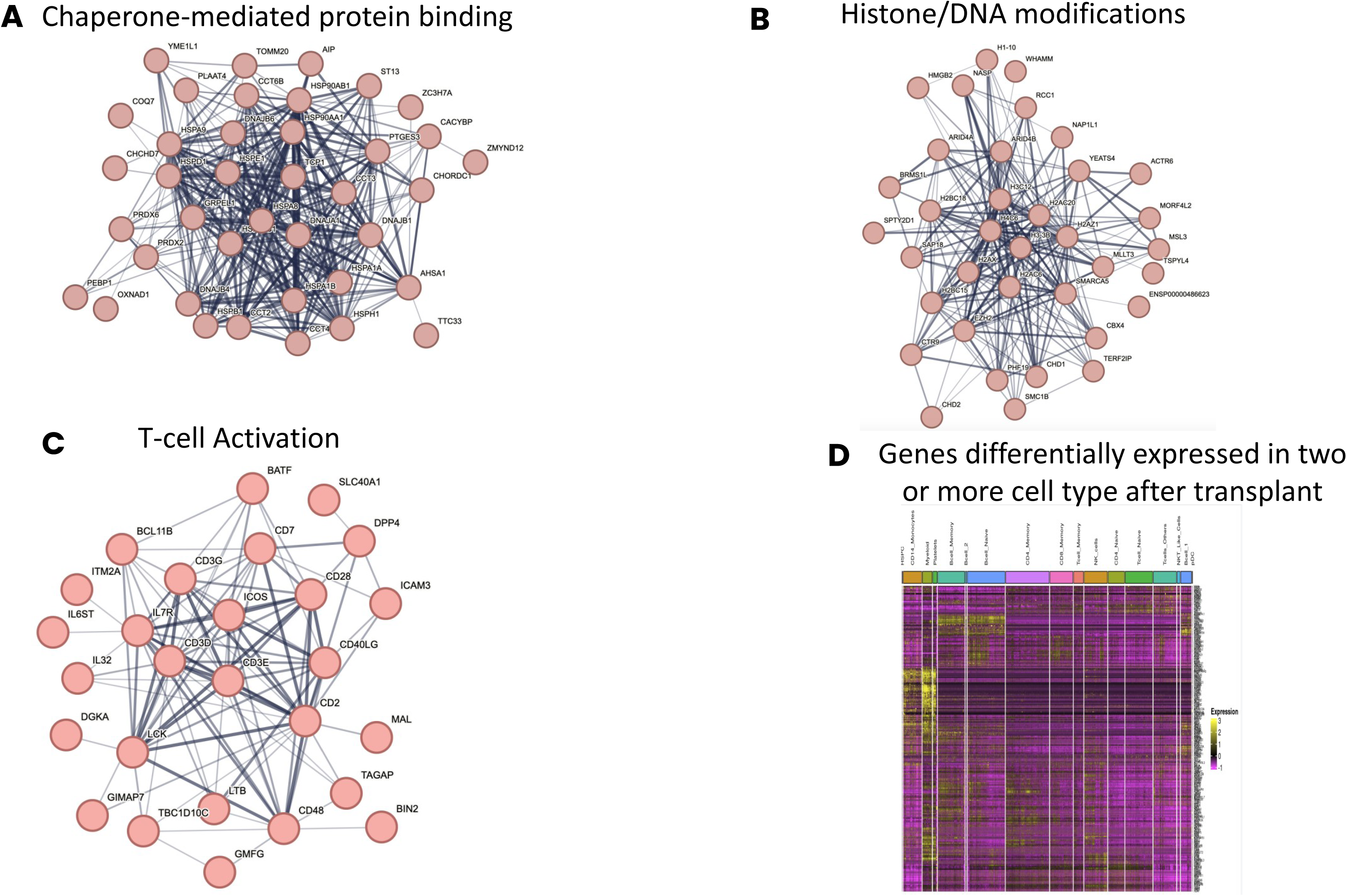
Major clusters of DEGs revealed by Markov clustering of DEGs in STRING in **A-C.** Pre-transplant CD8 memory cells, and post-transplant**. D.** Heatmap shows genes which were differentially expressed in two or more cells in post-transplant leukocyte subsets.

**Figure S2.**
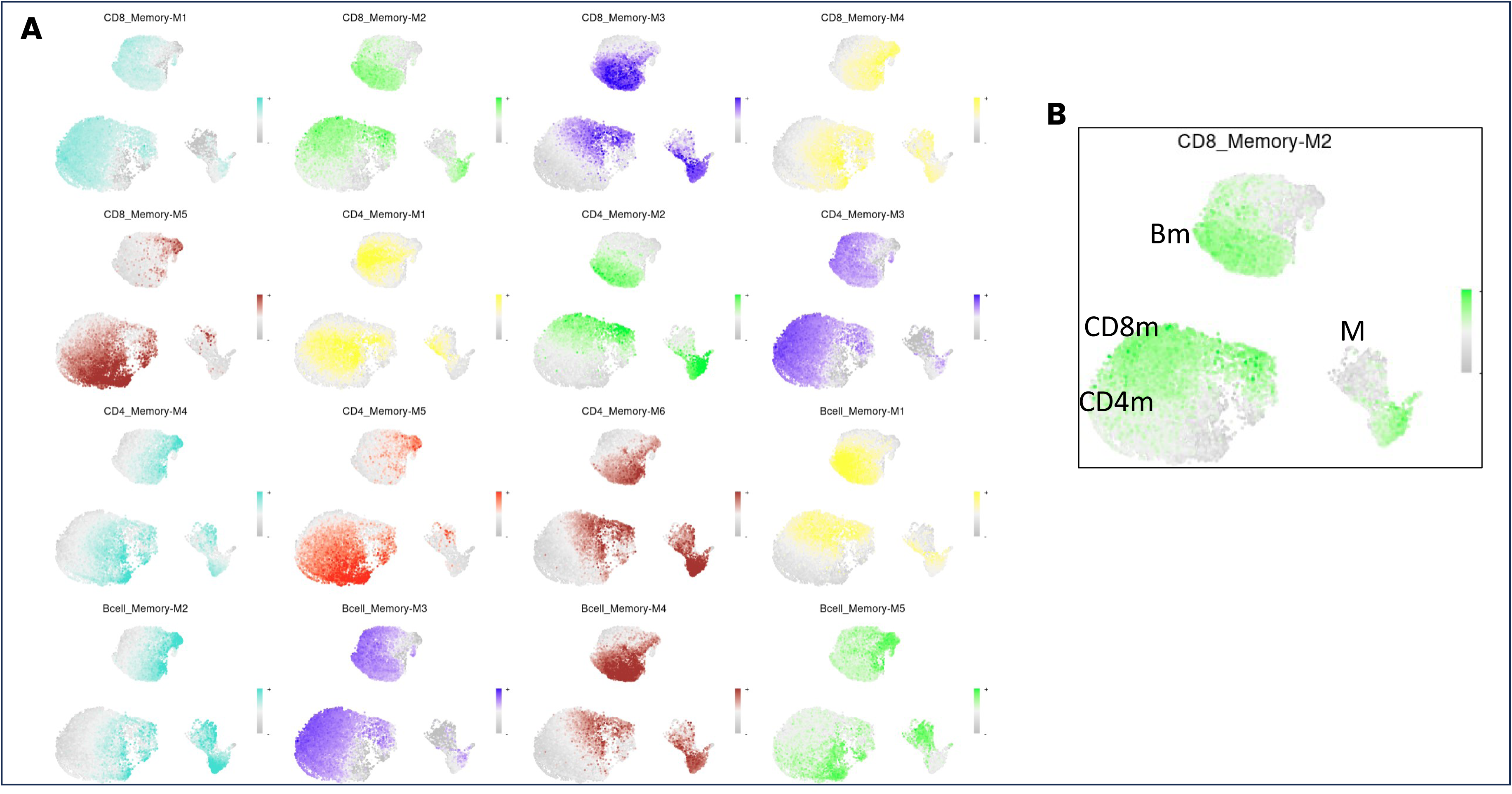
**A.** Panel of UMAPs shows cells in memory subsets of post-transplant CD8, CD4 and B-cells and myeloid cells which shows covarying expressing of genes in each module. **B.** UMAP shows location of memory (m) and myeloid (M) subsets.

**Figure S3.**
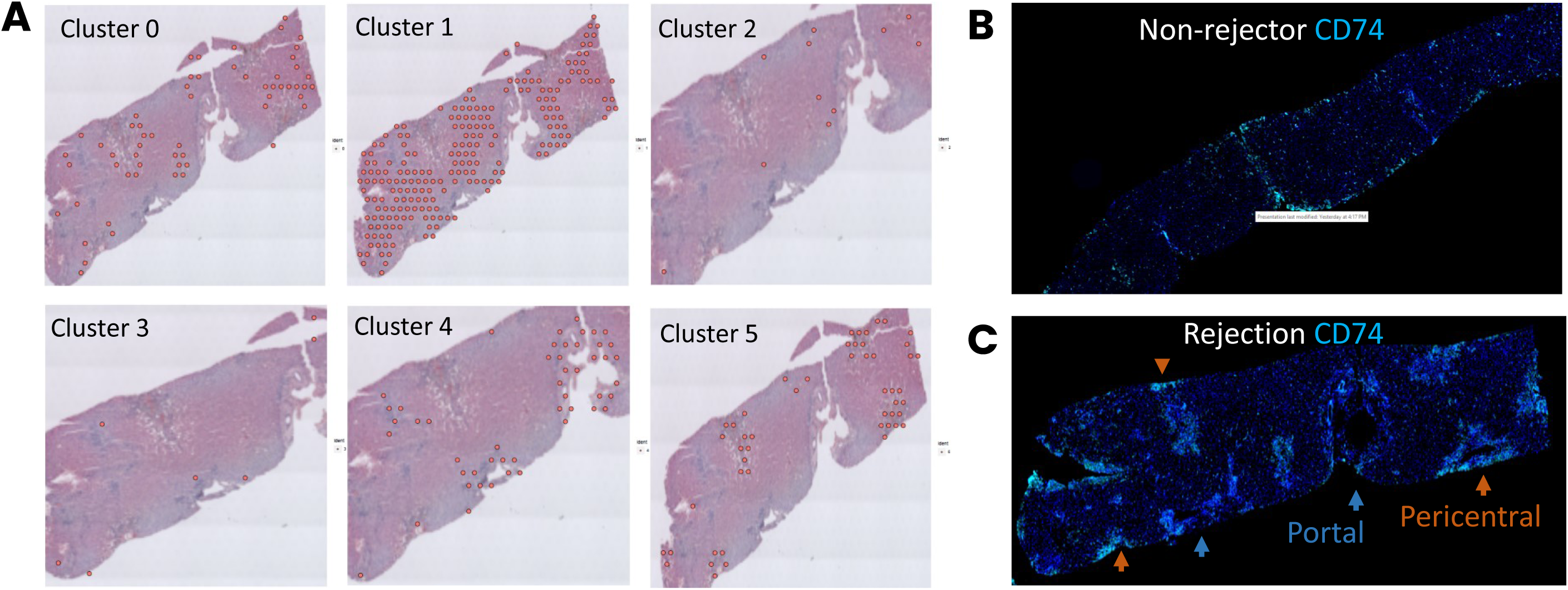
**A.** Location of six clusters identified in all biopsies, in the biopsy with rejection in Figure 3. Clusters 4 and 5 overlie the portal and pericentral regions, respectively. These six clusters were defined by the presence of unique anchor genes in spatial transcriptomes of all 7 biopsies. **B** and **C**. Immunostaining for CD74 (bright blue) in biopsies with no rejection (B) and rejection (C). Arrows point to portal (dark blue) and pericentral regions (rust red) that have been identified in Figure 5.

**Figure S4.**
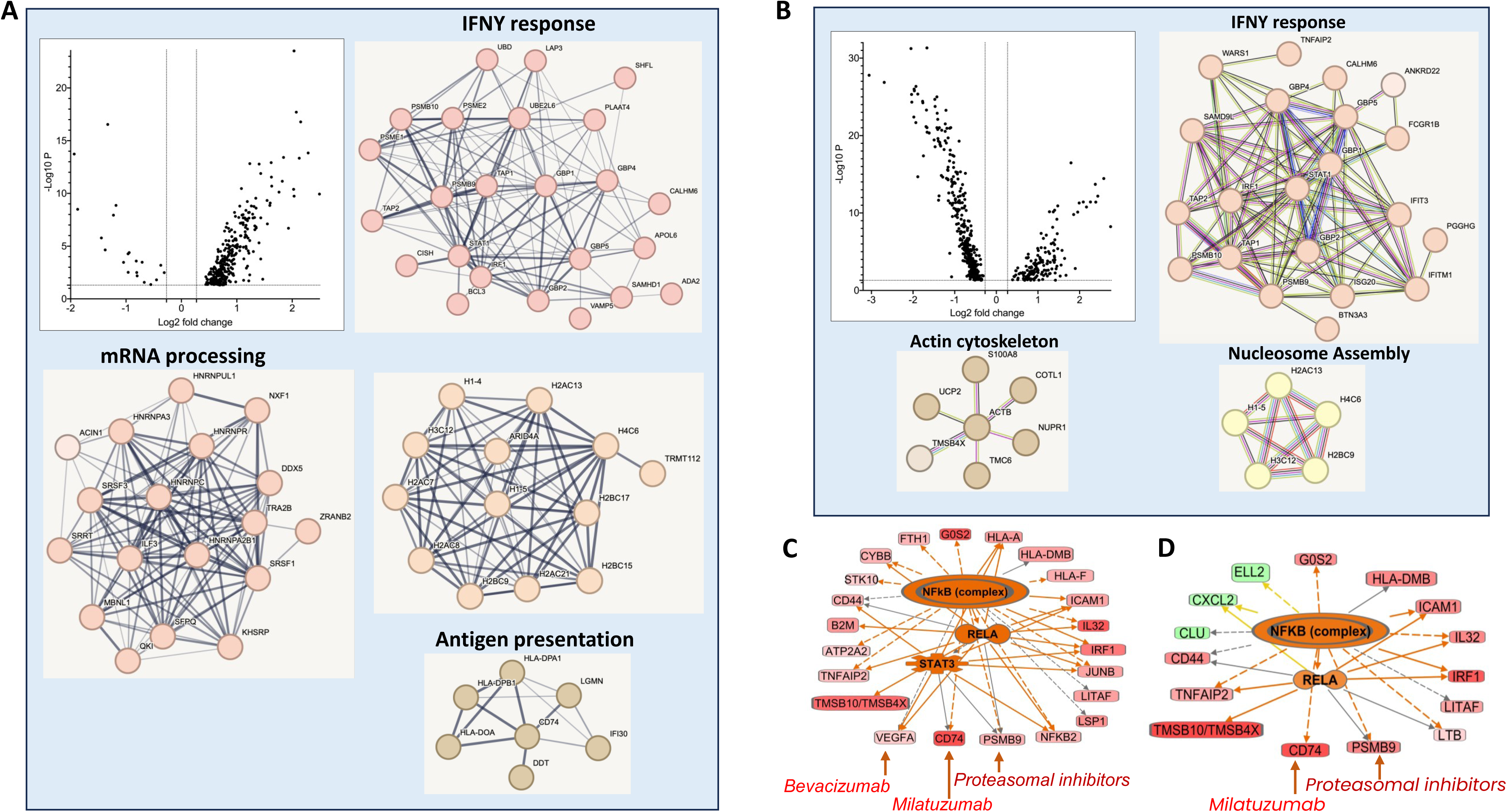
Volcano plots show all DEGs in **(A)** cluster 4 portal, and **(B)** cluster 5 pericentral regions of the allografts with rejection (R) compared with those without rejection (NR). Key genes are shown in volcano plots in Figure 3. Also shown in panels A and B are major functional clusters of upregulated DEGs. **C** and **D.** NFKB-regulated gene networks and drug targets within each network in portal (C) and pericentral (D) regions.

**Figure S5.**
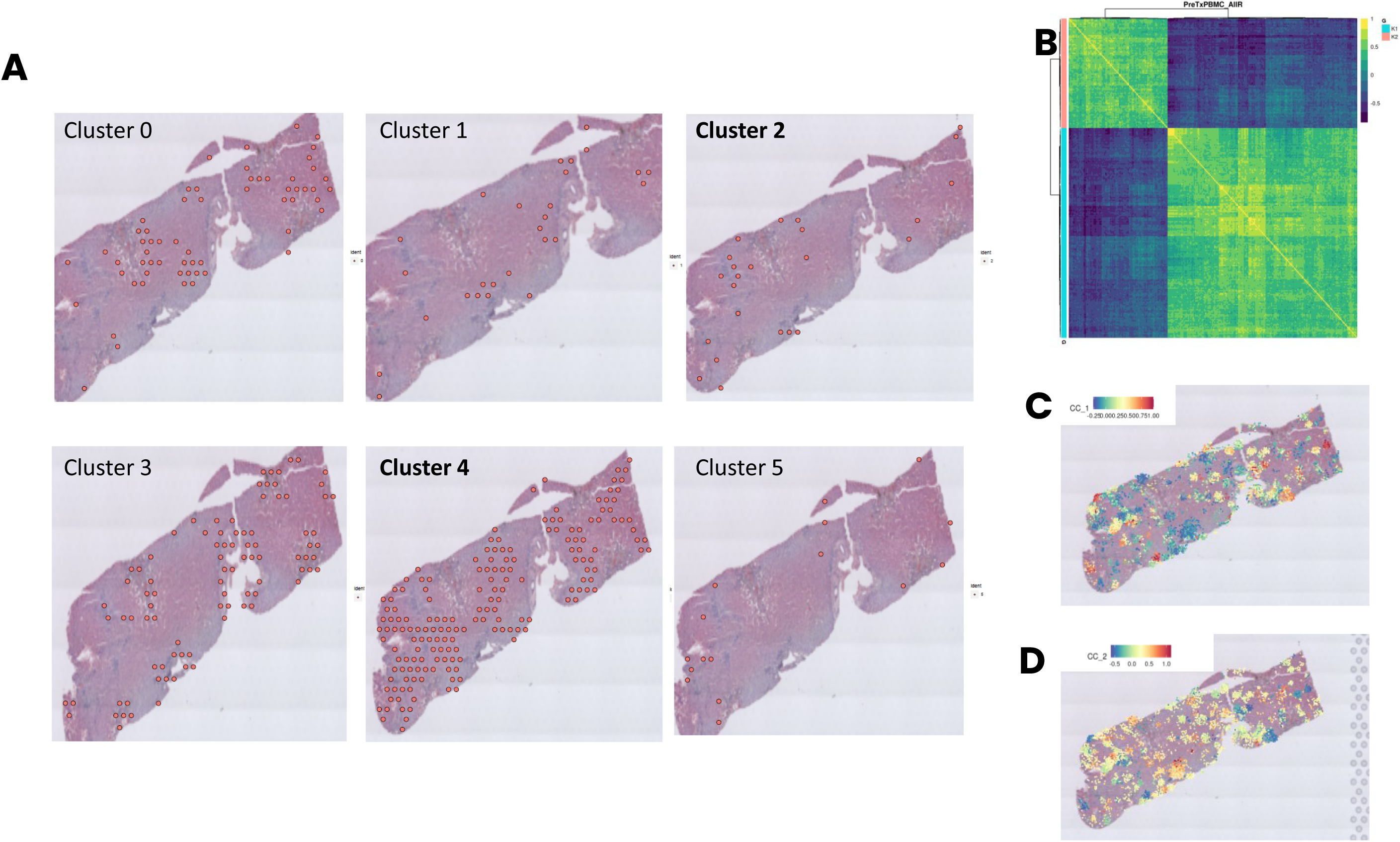
**A.** Location of six clusters identified in 5 biopsies with rejection, in the biopsy with rejection from Figure 3. Clusters 2 and 4 are dispersed in the intralobular region. **B.** Correlation analysis between DEGs in memory CD8, CD4 and B-cells in pre-transplant samples, with genes expressed in biopsy with ACR and cDSA. **C** and **D**. Location of the B-cell-rich cc1 and the T-cell-rich cc2 module in the biopsy with rejection.

**Figure S6.**
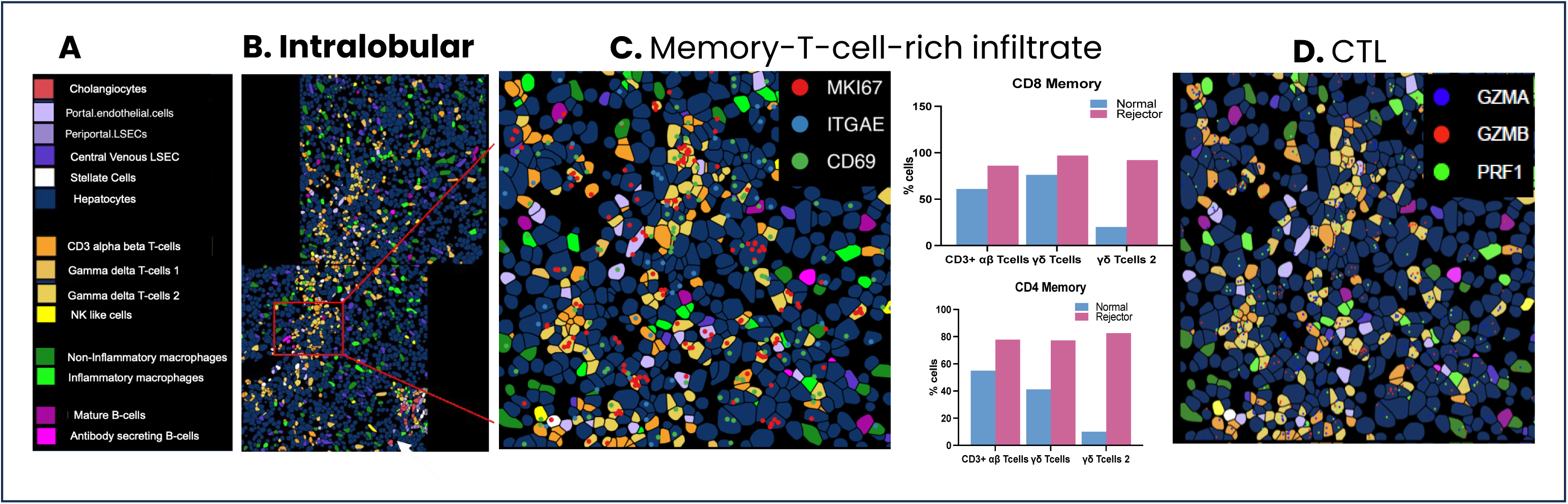
Spatial imaging of TCMR with cDSA. **A.** Color legend for cells in spatial images. **B.** Memory cell-rich infiltrate. Bar diagrams show alpha-beta and gamma-delta T-cells frequencies that express memory markers among CD8 and CD4 cells in the rejection biopsy and pre-transplant liver biopsy. **C.** Cytotoxic T-cells.

**Figure S7:**
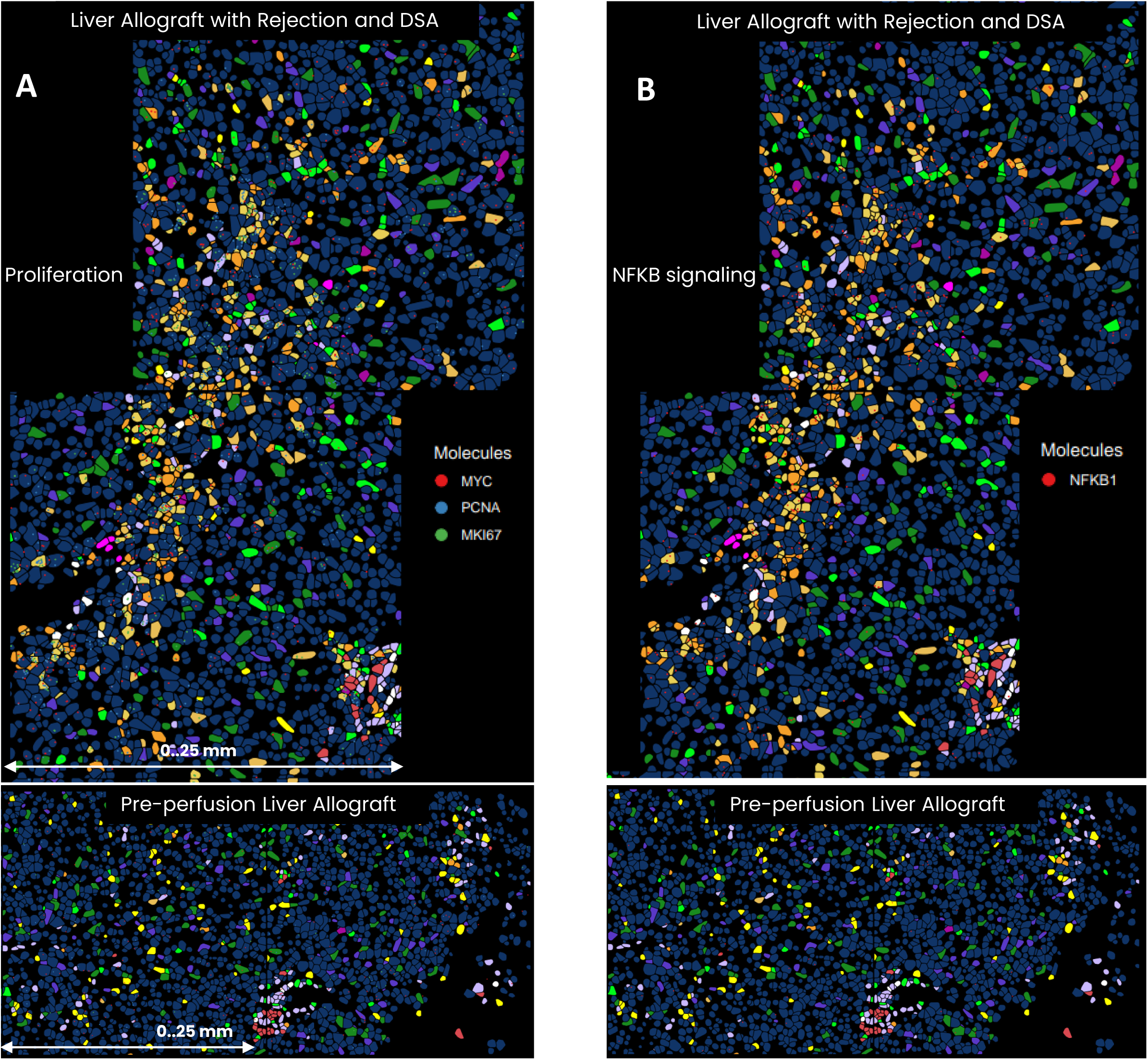
Additional germinal-center-like reaction genes for **A.** proliferation and **B.** NFKB signaling.

**Figure S8.**
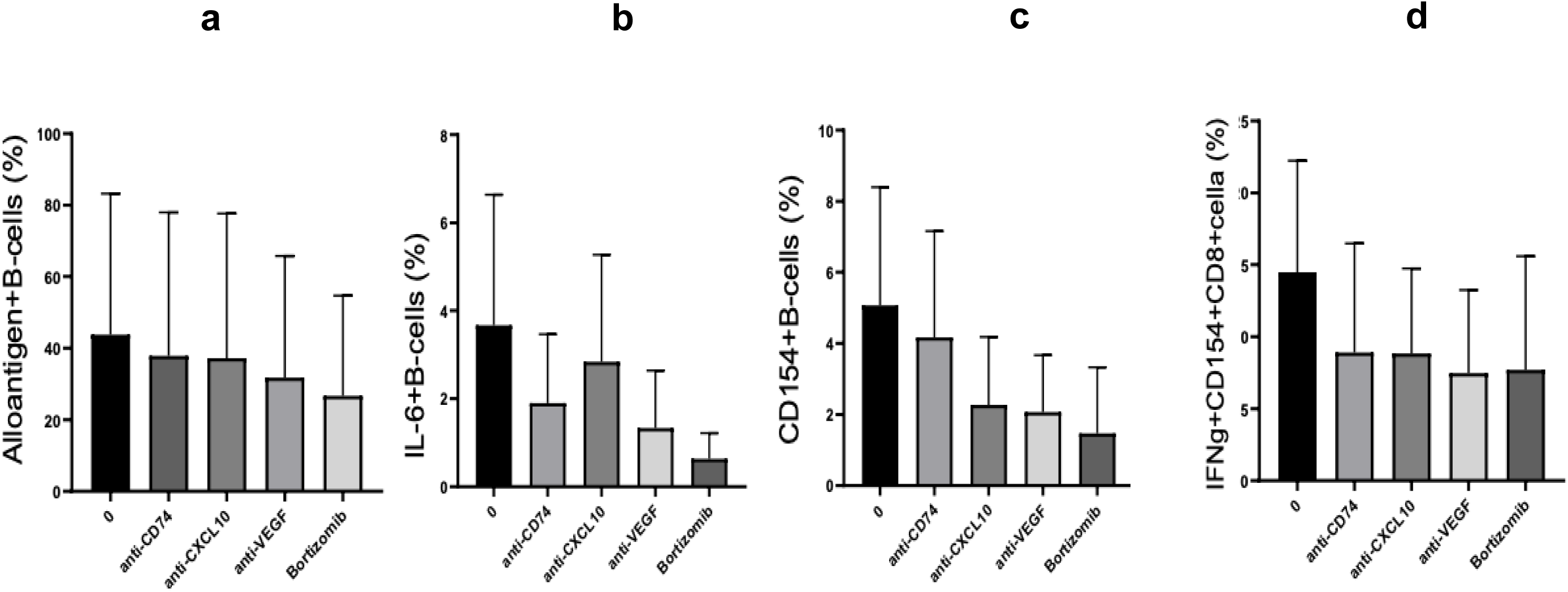
Effect of blocking novel drug targets on immune responses. The immune responses are measured as **a**) the frequency of B-cells that present alloantigen, **b**) allospecific IL6+B-cells, **c**) allospecific CD154+B-cells, and **d**) allospecific IFNγ+CD154+CD8. Each plot shows effects of blocking anti CD74, anti-CXCL10 (edelimumab), anti-VEGFA (bevacizumab), and bortezomib in co-cultures between HLA-mismatched alloantigen and responders from three pairs of responder and stimulator leukocytes.

**Figure S9.**
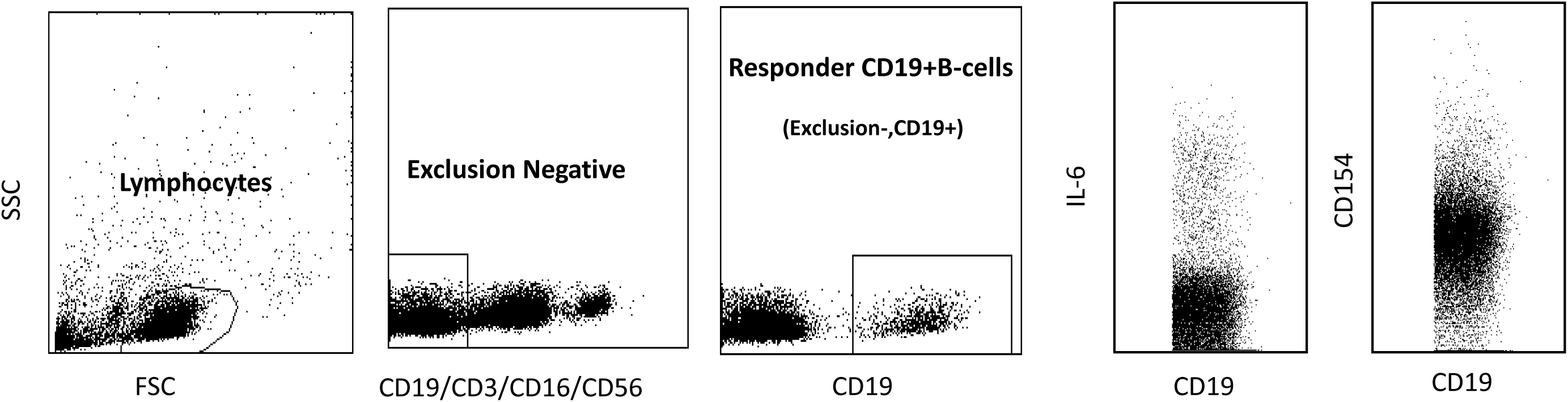
Total lymphocytes were gated based on Side Scatter (SSC) and Forward Scatter (FSC) gating, followed by selecting T-NK-NKT-negative cells (CD3/CD16/56 negative). Responder B-cells which are prelabelled with anti-CD19APCH7 are then separated out from Stimulator B-cells which are anti-CD19PECY7 labelled. Responder CD19+Bcells are then gated for anti-IL6PE and anti-CD154BV421 expression.

